# ESMBind and QBind: LoRA, QLoRA, and ESM-2 for Predicting Binding Sites and Post Translational Modification

**DOI:** 10.1101/2023.11.13.566930

**Authors:** Amelie Schreiber

**Affiliations:** Independent Researcher

## Abstract

In this study we discuss the viability of applying protein language models to the problem of predicting bindings sites of protein sequences from single sequences alone using Low Rank Adaptation (LoRA) and Quantized Low Rank Adaptation (QLoRA). No Multiple Sequence Alignment (MSA) or structural information for the proteins was used. Moreover, using LoRA and QLoRA shows improved performance over vanilla full finetuning, and significantly helps in mitigating overfitting. Also, due to the efficiency of LoRA and QLoRA, we are able to train the larger ESM-2 models on modest hardware, making the method very attractive and accessible. We also note that this technique serves as an important regularization technique and serves to improve generalization of models on unseen data.

## 1 Introduction

In this study we seek to determine to what extent protein language models alone can be applied to the task of predicting binding sites of proteins. It has been shown in recent studies that protein language models (pLMs) like ESM-2 may contain enough evolutionary and structural information to predict binding sites from sequence information alone. However, a crucial part of the method was scaling models up. In (GNH), the authors state “*We have shown that as more complex pLMs are employed to represent node features, the relative impact of the GNN architecture diminishes. This observation suggests that, to some extent, the structural information required for accurate binding site prediction is inherently captured by the pLMs themselves*.” Thus we seek to make finetuning of larger models more accessible to more people by implementing parameter efficient fine tuning strategies such as LoRA and QLoRA.

As pLMs become more user friendly and well documented, they become attractive options for binding site predictors as they allow one to use a single protein sequence, without multiple sequence alignment (MSA), and without any structural data for the protein. The prediction of binding sites can be approached as a basic binary token classification task, for which the model can be finetuned. In this study we discuss how to apply an important and simple regularization and parameter efficient finetuning strategy known as *Low Rank Adaptation* (LoRA), and its quantized version QLoRA. These strategies have been proven to provide better performance on various downstream tasks as compared to vanilla finetuning, and they prevent *“catastrophic forgetting”*. Moreover, their ability to help mitigate overfitting makes them an important regularization strategy for protein data, which is somewhat notorious for overfitting, especially when sequence similarity is not considered.1 We found that even the smallest ESM-2 model *esm2_t6_8M_UR50D* overfits early using vanilla finetuning. However, with the LoRA, the overfitting issue is mostly overcome with large enough datasets, and enough adapter layers with low rank, and it comes with the added benefit of being a more parameter efficient finetuning strategy, as well as showing improved performance in terms of the train/test metrics (accuracy, precision, recall, F1 score, AUC, and MCC).

### 1.1 Objective

The main objective here was to delve deeper into the performance metrics of a scaled-up model in predicting the binding and active sites solely based on single protein sequences, without MSA or structural data. Drawing from our past experiments with a smaller 8M parameter pLM, we were curious to see if a larger 35M parameter pLM could offer better predictive accuracy and reliability. We also plan to scale further to determine if performance continues to improve, and to discern if scaling laws with data such as with Chinchilla (Hoffman et. al.) holds. Determining the optimal size of the datasets based on model complexity will be part of this ongoing work. The other aspect of this project is making datasets, code, and models freely accessible and open to the community. Thus, we will be releasing datasets, the code to obtain those datasets, training scripts, code to evaluate the models, and many of the models themselves on Hugging Face to make them accessible to the community.

### 1.2 Methodology

We embarked on a meticulous training regime involving the ‘esm2_t6_8M_UR50D’ and ‘esm2_t12_35M_UR50D’, and ‘esm2_t33_650M_UR50D’. We chose these models due to their SOTA performance which is comparable to AlphaFold2 for the 3B parameter ESM-2 model, and due to their simplicity, requiring no MSAs or structural information. The backbone of this training were several datasets extracted from the UniProt database, housing around 209K, 550K, 600K, 770K, 1.1M, 2.6M, 11.4M, and 16M protein sequences annotated with binding site information and, where available, supplemented with active site data. A set of well-thought-out hyperparameters steered the LoRA and QLoRA adaptations, monitoring the performance carefully at different stages to glean a detailed understanding of the model’s evolving capabilities. We also tested models on publicly available data from the PDB available at (GNH), and found the performance on these datasets to be poor, although some of the metrics imply good generalizability of the smaller models we’ve trained so far. Explaining this performance drop is an as yet unsolved problem which we have only conjectures for. The issue could be in the difference in annotations between UniProt and the PDB.

It could also be due to discrepancy in dataset size, as these datasets are quite small (approximately 5K proteins). This could also be due to the fact that we have not scaled up the datasets and model sizes yet, and what little scaling we have done has not been in a 1-to-1 fashion as is recommended in (Hoffman et. al.). Yet another explanation could be that the train/test splits in (GNH) are random, going against the advice of (WPT). Only proteins with binding site annotations were selected in all datasets. For all of the datasets, the proteins were sorted by UniProt family and then random families were selected to be separated out for the train/test split, which was approximately an 80/20 split. A small subset of the sequences (approximately 50) were excluded due to having only partial binding site annotations. Active sites were also merged with binding sites. The sequences longer than 1000 residues long where then segmented into non-overlapping chunks of 1000 amino acids or less to account for the 1022 sized context window of the base model. Class weights were used due to the imbalance between binding sites and non-binding sites. However, this may be changed later on to only be used during early stages of finetuning or perhaps removed altogether.

### 1.3 Findings

As we navigated through the results, we noticed an encouraging trajectory. The model not only exhibited an uptick in key metrics such as accuracy, precision, recall, F1 score, AUC, and MCC, but also hinted at areas where we could fine-tune to achieve even better results. We noticed that the 209K dataset overfit after the third epoch with ‘esm2_t6_8M_UR50D’, and the 550K dataset showed little overfitting, but also little performance improvement after 6 epochs. We then decided to scale to the larger model, ‘esm2_t12_35M_UR50D’, which overfit early on the 550K dataset, showed some improvement on the 770K dataset but still overfit, and results for the 1111K dataset showed significantly less overfitting, especially with more QLoRA adapter layers. Testing with the 2600K dataset are still in progress. While LoRA and QLoRA show dramatic improvements in regards to overfitting, and noticible performance improvements in terms of test metrics due to this, overfitting and obtaining good generalization are still challenges to be overcome by finding the correct dataset size and scaling the data in a 1-to-1 fashion. We also note that adapting more weight matrices with LoRA or QLoRA provides additional reduction in overfitting on smaller datasets that were note scaled in a 1-to-1 fashion. In other words, adapting more weight matrices with LoRA and QLoRA serves as a very effective regularization technique. We found that models with all weight matrices adapted by LoRA or QLoRA had less overfitting on small datasets than models only using query, key, and value weight matrix adapters. Moreover, we are finding that QLoRA has improved additional regularization effects, reducing overfitting even more than LoRA. We also found that performance is not significantly improved by increasing the rank of the LoRA or QLoRA, and that a lower rank of 2 is sufficient for this task. Determining how rank effects overfitting is part of our ongoing work.

## 2 LoRA and QLoRA Helps Prevent Overfitting

LoRA and QLoRA have been shown to be an important and successful regularization technique and for some tasks can provide improved performance over standard full parameter finetuning. We find that this holds true for our task of predicting binding sites and that overfitting is dramatically improved with LoRA, and even moreso with QLoRA. We found that lower rank of the LoRA provide better performance both in terms of overfitting and in terms of metrics on the test data. Since QLoRA allows for training billion parameter models on (top of the line) consumer grade hardware2, it is likely that we will be able to finetune the 650M and 3B parameter ESM-2 model without issue, albeit with small batch sizes. The more pressing issue is dataset size, and if (Hoffman et. al.) scaling laws hold for ESM-2, we may run out of data at the 650M parameter model. Below, in Table 1 we see an example of how adding more QLoRA adapters induces more regularization and helps prevent overfitting. In this table, we have the train and test metrics for two QLoRA adapters for ‘esm2_t12_35M_UR50D’, with only the query, key and value weight matrices adapted, and with all weight matrices adapted.

**Table 1:**
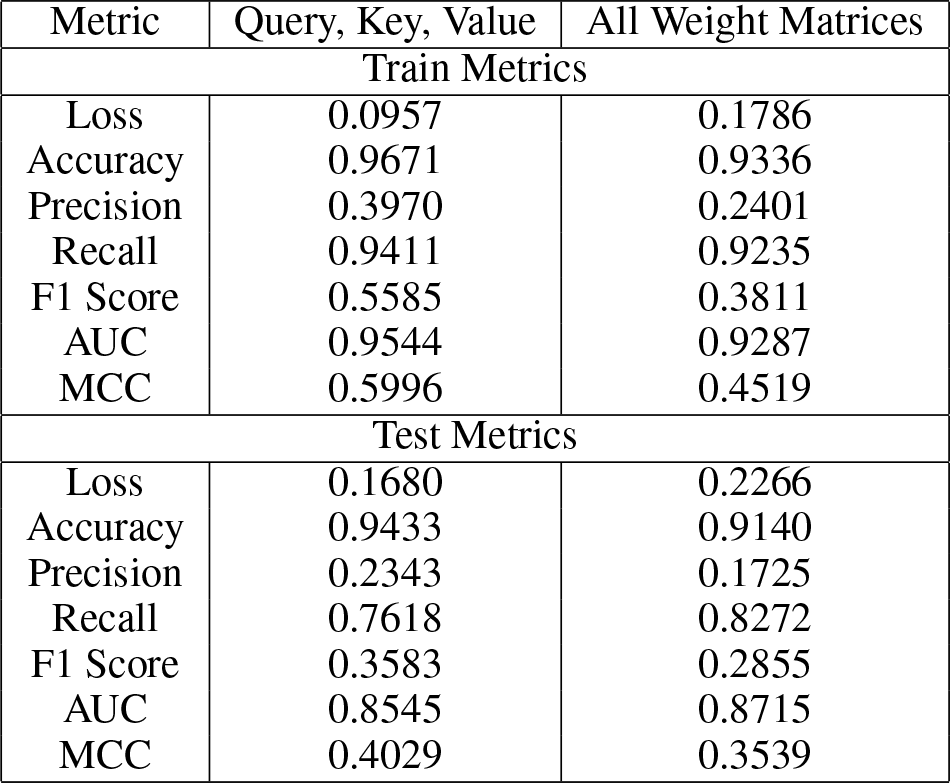
Performance Metrics for Different Weight Matrices.

Determining whether or not these methods are viable as an alternative to structural models and models that require MSA will be the main goal of this project. So far results have been promising on our train test splits. However, when testing the models on the relatively small datasets provided in (GNH), we see a large performance drop for the 35M parameter model, which has yet to be explained, but is likely due to not following (Hoffman et. al.) and perhaps due to the random train/test splits used in (GNH). Our sequences in the train/test splits were split by family to avoid sequence similarity. We intend to do further testing with BLAST and perhaps will use UniRef to split the datasets in the future. Another possibility is a potential difference in annotations between the UniProt data that we used, and the PDB datasets used in (GNH).

### 2.1 Models and Training

We finetuned several of the ESM-2 models with LoRA and QLoRA, beginning with the 8M parameter model, then 35M, 150M, and 650M. Most models overfit prior to completing 3 epochs, and none were trained for more than 4 epochs. We also trained a small 8M parameter model on the 600K dataset to compare and show that full finetuning, even with aggressive dropout, overfits early and much more severely than LoRA and QLoRA finetuning, as can be seen in the Table 2 below.

**Table 2:** Full Finetuning Overfitting.

| Metric | Train | Test |
| --- | --- | --- |
| Loss | 0.1365 | 0.2910 |
| Accuracy | 0.9656 | 0.9233 |
| Precision | 0.3862 | 0.1489 |
| Recall | 0.9618 | 0.5335 |
| F1 Score | 0.5511 | 0.2328 |
| AUC | 0.9638 | 0.7327 |
| MCC | 0.5978 | 0.2533 |

Comparing three of the models of the same size (8M) trained on the 600K dataset in a table showing the differences between the train and test metrics, we get the following Table 3:

**Table 3:** Train/Test Metric Differences.

| Metric | Full Finetuning | Query, Key, Value | All Weight Matrices |
| --- | --- | --- | --- |
| Loss | -0.1545 | -0.0723 | -0.0480 |
| Accuracy | 0.0423 | 0.0238 | 0.0196 |
| Precision | 0.2373 | 0.1627 | 0.0676 |
| Recall | 0.4283 | 0.1793 | 0.0963 |
| F1 Score | 0.3183 | 0.2002 | 0.0956 |
| AUC | 0.2311 | 0.0999 | 0.0572 |
| MCC | 0.3445 | 0.1967 | 0.0980 |

For LoRA and QLoRA, various hyperparameters were adjusted to help with overfitting, and we began with using Weights and Biases sweeps for the smallest 8M parameter model, but this was infeasable for larger models due to hardware constraints. The hyperparameter configurations are released with the models on Hugging Face, including the LoRA and QLoRA configurations. Most models were trained with a LoRA rank of 2 as sweeps for the 8M parameter model did not indicate this effecting performance very much, and reducing rank seems to help with overfitting to some small degree, implying binding site information is likely located in lower intrinsic dimension (see (Hu et. al.) and (AZG)). Various LoRA weight matrix configurations were used, until training larger 150M parameter models and it was found that more weight matrices being adapted by LoRA (or QLoRA) provided more regularization and reduced overfitting significantly more than using only query, key, and value adapters. We also found that full finetuning overfit very early and drastically more than with LoRA and QLoRA finetuning. Part of this might be explained by research presented in (RC). The training metrics for the models can all be found on Hugging Face.

### 2.2 Hardware

All 8M parameter models were trained on a single 24GB A10 GPU using Lambda Labs, or two smaller 8GB GPUs locally. Some of the 35M parameter models were also trained on the 24GB A10 GPU, and some were trained on a single H100 80GB GPU. The 650M parameter model was trained only on the H100 to allow for large batch sizes. This along with QLoRA allowed for very large batch sizes, as QLoRA trained less than 1% the parameters compared to the trainable parameters in full finetuning. We note, it is possible to train the 150M, 650M, and 3B models on the 24GB A10 GPU.

## 3 Limitations and Future Improvements

While the 650M parameter model was trained on approximately 12M protein sequences with binding site annotations from UniProt, it could easily have been trained on the full 16M, and this is one planned improvement we intend to make given the compute is available. We also note that these models can provide useful embeddings for other models, and having been finetuned for predicting binding sites, will likely improve performance of structural models more than the embeddings from the pretrained ESM-2 models or other protein language models which have not been finetuned for predicting binding sites. Further comparison of the method could also be done, however, comparing to other models that predict binding sites is difficult due to widespread data leakage in those models, where things like sequence similarity are not considered. A simple random train test split, or even more complicated splits that don’t take sequence similarity into account cause overfitting to occur in models without showing any of the traditional signs of overfitting, thus making a comparison to other methods extremely difficult. Moreover, comparing structural models to sequence only models is difficult due to the compute necessary to predict the 3D folds of proteins first. Given the time and compute necessary for such a comparison, and assuming structural models are trained with structural and sequence similarity in mind, such a comparison could be made for standard models like fpocket, p2rank, PointSite, etc. and this is one way we could improve the current work. We also note that training the larger ESM-2 models such as the 3B and 15B parameter models could feasibly be done with very modest hardware requirements, and this is indeed an improvement we would like to make assuming the compute to do so is available. Another improvement that could be made is a more extensive hyperparameter search for smaller models, then using *μ*-Transfer to transfer hyperparameters to larger models, a more optimal set of hyperparameters could be found. Additionally, more extensive data on how LoRA and QLoRA improve generalization and reduce overfitting could be done by performing full finetuning for larger models, then applying LoRA or QLoRA to progressively larger subsets of the weight matrices could be done to show how the overfitting is reduced, and to what degree, by the various weight matrices. This might also be an interesting direction for future projects related to predicting other properties of proteins such as post translational modifications or function prediction, which is already underway.

### 3.1 Conclusion

LoRA and QLoRA have shown improved performance metrics compared to full finetuning, as well as dramatic improvements in overfitting when adding in more adapter layers. Additionally, sorting the train/test split based on protein families has helped mitigate some of the overfitting as well. Despite this, overfitting is still a challenge, and further techniques to mitigate this are required. Additionally, testing the models on the data available in (GNH) indicates potential issues. In particular, we noticed that (GNH) uses random train/test splits, going against the recommendations of (WPT), and contrary to our approach of separating out test data by protein family to help avoid sequence similarity between the training data and the test data (which could still be improved). The overfitting issues also made it obvious that there was an issue with the way we were scaling the datasets initially, that is, in not yet following the recommendations of (Hoffman et. al.).

However, it also prompted us to expirement with which weight matrices we were applying the LoRA and QLoRA adapters to, at which point we discovered that more weight matrices adapted by LoRA or QLoRA provided additional regularization. We also found that QLoRA was even more effective at regularization than LoRA, which already reduced overfitting drastically. Furthermore, we believe that improving precision at the expense of other metrics is also a potential point where the model could be improved, as this may help mitigate costs of testing in a lab, due to reduced false positives, if scaling up provides more performance improvements as suggested in (GNH). It’s worth noting that the presence of many under-annotated binding sites in UniProt datasets might lead our model to identify numerous true binding sites but also predict many false positives, resulting in high recall but low precision. To help, we are also taking inspiration from (HYMWL) and building ensemble models. More work is needed to apply this method effectively to predicting binding sites of protein sequences, but the fact that it requires no Multiple Sequence Alignment (MSA), no structural data for the proteins, the simplicity of the method, the promising metrics on the UniProt datasets which appear comparable to SOTA structural models, as well as the ability to finetune larger ESM-2 models on modest hardware make it very attractive. Code and models are being released as the project progresses.

## 4 Supplementary Material

### 4.1 Post Translational Modification

We also finetuned three different sizes of ESM-2 (8M, 35M, and 150M) for predicting post translational modification sites of protein sequences using standard finetuning, LoRA, and QLoRA. Similar to the task of predicting binding sites, we treat the problem as a binary token classification task. Below in Table 4 and Table 5 are the metrics for the models we have trained and computed metrics for thus far.

**Table 4:**
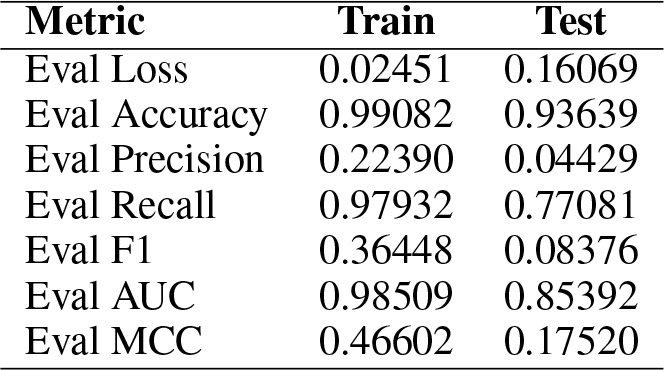
Comparison of Train and Test Metrics for ESM-2 (8M) LoRA.

**Table 5:**
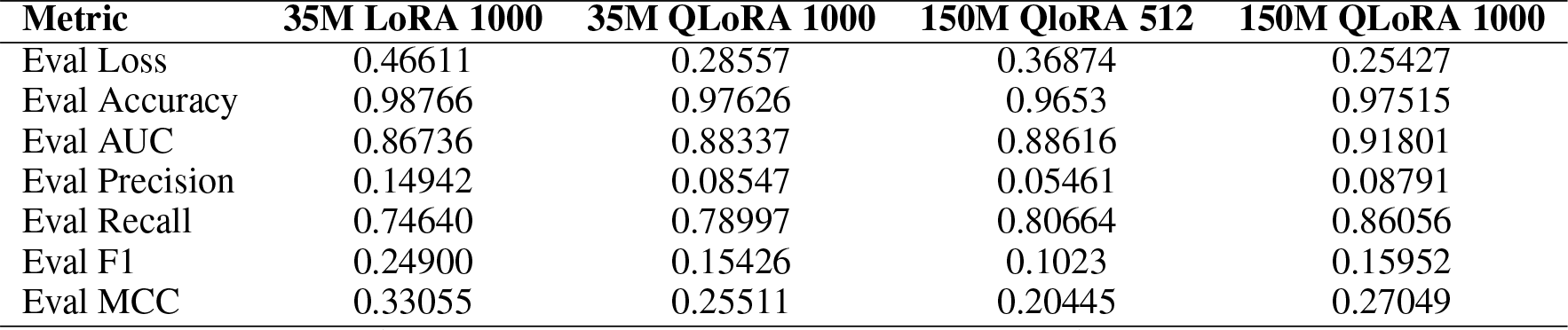
Comparison of Metrics Across Four Models.

Interestingly, we note that a longer context window of 1000, as apposed to the 512 context window, seems yield improved performance. The ESM-2 8M parameter model was trained on approximately 500K protein sequences with post translational modification site annotations from UniProt. The 80/20 train/test split was created based on protein family to account for protein homology. Similarly the 35M parameter models, and the 150M parameter models were trained on approximately 2.1M protein sequences with PTM site annotations from UniProt, with an 80/20 train/test split based on protein families to account for sequence similarity between the train and test datasets. For finetuning, we used class weights due to the significant imbalance in the data, with PTM sites occuring very infrequently compared to non-PTM sites. We note that there are approximately 2M protein sequences in UniProt currently with post translational modification site annotations, but without protein family annotations, which the model can be tested on for generalization capabilities, and this is in progress. Obtaining comparisons of the train and test metrics to test for overfitting is also in progress. The best performing 150M model will also be available at https://neurosnap.ai/services for use without the need for code.

### 4.2 Data, Code, and Model Availability

We are making all code used to preprocess the data, as well as training scripts, and models available on Hugging Face. We have created two “Hugging Face Collections”. One for LoRA models, and one for the QLoRA models.

All LoRA models for binding site prediction can be found at: https://huggingface.co/collections/AmelieSchreiber/esmbind-esmb-for-protein-binding-sites-64f9e4c3f210b74ac92b8b55.

All QLoRA models for binding site prediction can be found at: https://huggingface.co/collections/AmelieSchreiber/qbind-qlora-for-esm-2-binding-sites-prediction-651612ea01d873dd19124bfa.

All models for post translational modifications can be found at: https://huggingface.co/collections/AmelieSchreiber/esm-ptm-esm-2-for-predicting-ptm-65206f49706c75514869aa96. See also https://neurosnap.ai/services to use the best performing 150M parameter model without the need for code.

### 4.3 What is… a LoRA?

In the realm of deep learning, the concept of Low Rank Adaptations (LoRAs) was first introduced by (Hu et. al.). These LoRAs provide an efficient alternative to the traditional fine-tuning of neural networks. The process begins by freezing the pre-existing weights of a layer in the neural network. For instance, in the context of a transformer’s attention mechanism, this could involve freezing the weights of the query, key, or value matrices—often represented as *W*_*Q*_, *W*_*K*_, and *W*_*V*_. Following this, a LoRA layer is introduced to one or more of these pre-trained weight matrices. If we consider *W* to be a frozen weight matrix, the LoRA layer would take the form of *W* + Δ*W*, wherein Δ*W* = *BA* constitutes the LoRA. Typically, these are low-rank decompositions, with *A* ∈ R^*r*×*d*^_*in*_ and *B* ∈R^*d*^_*out*_ ×*r*, where the original weight matrix is *W*∈ R^*d*^_*out*_ ×*d*_*in*_. It is common for *r* to be significantly less than min {*d*_*in*_, *d*_*out*_}. The application of LoRAs only provides significant benefits when *r* is much smaller than the input and output dimension. Nevertheless, we can still opt for a smaller *r* and implement a LoRA in lieu of conventional fine-tuning. Empirical evidence suggests that in many cases, selecting *r* = 4 or *r* = 8 is sufficient—even for large weight matrices such as the query, key, and value matrices of a transformer’s attention mechanism. Let’s now explore a scenario where the application of a LoRA does not yield any substantial benefits in terms of reducing the number of parameters:

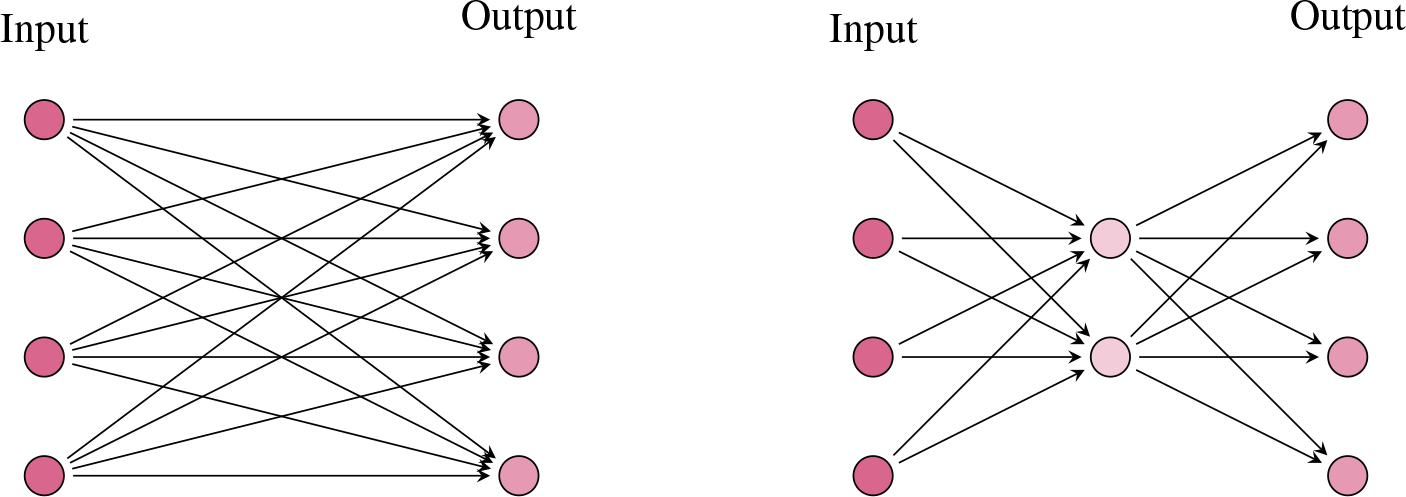

Here, we see that the number of parameters for the LoRA layer Δ*W* = *BA* is the same as the original layer *W*, where we have 4× 2× 2 = 16 parameters for the LoRA (on the right), and 4 × 4 = 16 parameters for the original frozen weight matrix (on the left). Next, let’s look at an example that gives us 40% the parameters of the frozen weight matrix:

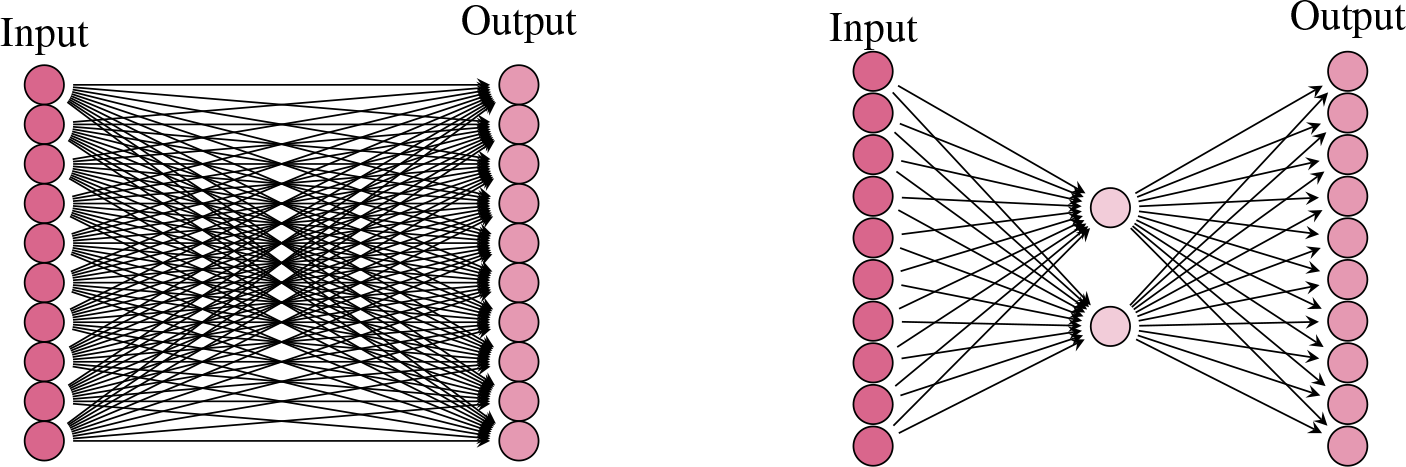

Here we see the original (frozen) weight matrix has 10^2^ parameters, and the LoRA has only 10 × 2 × 2 = 40 parameters. In most cases, we have that the rank (this is the number of neurons in the middle layer of the LoRA) of the frozen matrix is much smaller than the input and output dimensions, and there is in fact a drastic reduction in parameter count. As an example, we might have an input and output dimension of say 100, in which case the weight matrix has 100^2^ = 10, 000 parameters. However, the rank of this matrix is very often much lower than 100. In practice, it was shown that choosing *r* = 4 for the query, key, and value matrices is often more than sufficient for a LoRA as the middle dimension. In this case, we would get 100 × 4 × 2 = 800 parameters in the LoRA, which is less than one tenth the original parameter count. Once we have such a LoRA in place, we can train it on some downstream task, and then add the LoRA weight matrix *BA* to the original (frozen) weight matrix *W* to obtain a model that performs well on this new task. Importantly, LoRAs can help with issues such as overfitting, which can be a significant issue when learning on protein sequences. This, along with the parameter efficiency and a need to train larger models is why we decided to adopt LoRA as a finetuning strategy. Moreover, the simplicity of using a LoRA for parameter efficient fine tuning using the Hugging Face PEFT library makes it an attractive option. It also became clear early on that performance can actually *increase* with the use of LoRA and QLoRA, thus providing further motivation to adopt it as a strategy.

### 4.4 What is… a QLoRA?

In (DPHZ), the concept of QLoRA was introduced. In the proposed method, QLoRA (Quantized Low Rank Adapters), a streamlined finetuning approach is presented to significantly reduce memory consumption, making it feasible to finetune large language models (LLMs) with up to 65 billion parameters on a single 48GB GPU, while maintaining the model’s full 16-bit task performance. The core innovation lies in backpropagating gradients through a 4-bit quantized, frozen pretrained language model into Low Rank Adapters (LoRA). This process is facilitated by several memory-saving innovations: (a) the introduction of a new data type, 4-bit NormalFloat (NF4), optimized for normally distributed weights, (b) Double Quantization, which further reduces memory footprint by quantizing the quantization constants, and (c) Paged Optimizers to manage memory spikes during training. Through QLoRA, the Guanaco model family was trained, achieving superior performance on the Vicuna benchmark, outperforming previous models while drastically cutting down the finetuning time to 24 hours on a single GPU.

The fine-tuning process in QLoRA is encapsulated in a succinct formula which integrates the quantized weights and Low Rank Adapters (LoRA) in a single linear layer computation. This formula, *Y*_BF16_ = *X*_BF16_ doubleDequant 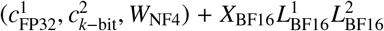, elegantly combines the input tensor, quantized weights, and LoRA in computing the output tensor, with gradients only required for the adapter weights, not for the 4-bit weights. Through these innovations, QLoRA not only significantly diminishes the memory footprint required for fine-tuning but also makes it feasible to fine-tune huge models on modest hardware, marking a significant stride in memory-efficient fine-tuning of large language models.

### 4.5 Why Do LoRA and QLoRA Help Improve Generalization?

The theoretical underpinnings of finetuning with LoRA and QLoRA were in part inspired by the work (AZG). This work connects generalization capabilities of models to the intrinsic dimension of data. We also not several recent works relating geometric compression and information theoretic compression (CKB), and studying how the intrinsic dimension of data evolves through the layers of the ESM-2 protein language models (VCAC), (VDCLAC). We conjecture that this is the source of the regularization and improved generalization to unseen data when using LoRA and QLoRA. In particular, when a model is inclined to overfit, we can force more geometric compression onto the model by choosing a rank for our LoRA or QLoRA that is lower than the intrinsic dimension of the data in any particular layer. We hypothesize that choosing a rank much larger than the intrinsic dimension of the embeddings in any given layer provides an over-parameterized LoRA or QLoRA and that it is sufficient to choose a rank approximately equal to the average (or perhaps the maximum) intrinsic dimension of the training data’s embeddings. We also theorize that a curriculum learning strategy might be developed based on the intrinsic dimension of the protein embeddings, whereby less complex proteins with lower intrinsic dimension are trained on first. We leave testing these theories to future work.

### 4.6 Why a Sequence Based Transformer Model?

There are two major classes of binding site prediction models, sequence based models, and structural based models. While it is often believed that structural models perform best, proteins with predicted 3D folds and structures represent a tiny amount of the proteins available in various databases. Moreover, although models such as ESMFold provide atomically accurate prediction of protein structure 60 times faster than that of AlphaFold2, the computation of 3D structures is still a bottleneck and most proteins remain without predicted structures, making structural models all but useless without the compute to predict the structures for all proteins of interest. Moreover, this limits the training data volume of structural models. Add to this the fact that the attention maps of transformer based protein language models recapitulate the contact maps of proteins, and the fact that ESMFold is essentially *“just”* a protein language model with a folding head on top, we see that protein language models, when scaled up, do in fact encode enough information to predict binding sites of proteins without any structural information, using only the protein sequence.

Additionally, due to the fact that no multiple sequence alignment is required, they become less cumbersome to use, and the barrier to entry becomes lower making these models more accessible to a wider audience. We also note the relative ease in applying LoRA and QLoRA to ESM-2 using Hugging Face’s Parameter Efficient Fine Tuning (PEFT) library. This makes even the larger ESM-2 models accessible and means that more people can finetune these models on relatively modest hardware, democratizing AI for studying proteins. So, with all of this in mind, we chose to adopt the transformer based ESM-2 models as our base model, and chose to adopt PEFT finetuning strategies such as LoRA and QLoRA. We also note, for those interested in structural models, that the embeddings of the LoRA and QLoRA finetuned models can be subsequently used in structural based geometric deep learning models to enhance their performance similar to what is presented in (WRXL). With the LoRA or QLoRA finetuned embeddings, the performance improvement of the structural models is likely to be enhanced even further. Thus we advocate for better sequence based methods such as this one, which can augment and improve structural models.

### 4.7 Data Processing

Our datasets were created by first searching UniProt for sequences that had binding site annotations, of which there were approximately 16.7M. A TSV file with columns “Protein families”, “Binding sites”, “Active sites”, and “Sequence” was downloaded from UniProt. Next, the sequences were sorted by family, and random families were selected to be separated out for test data to prevent data leakage, creating either an 80/20 or 70/30 train/test split. Next, any sequences with incomplete binding site annotations were discarded, removing 37 protein sequences in total. We then merged all active site annotations with binding site annotation, and converted this to binary labels with zeros for non-binding sites, and ones for binding sites. We then split all sequences longer than 1000 residues into non-overlapping chunks of length 1000 or less.

Similarly, the corresponding merged binary labels were also split. Next, a percentage of the sequences in the training data and test data were removed at random to make the datasets smaller for various model sizes. We created datasets with 550K, 600K, 770K, 1.1M, and 2.6M to use for the smaller 8M and 35M parameter ESM-2 models. We also created a dataset of 12M sequences to use with the 150M parameter ESM-2 model, and used the full 16M dataset for the 650M parameter ESM-2 model. We note, the 16M dataset is somewhat small for the 650M parameter model according to (Hoffman et. al.), where a 20-to-1 ratio of tokens to model parameters is suggested, and scaling is to be done in a 1-to-1 fashion, that is, doubling the models size implies one should double the dataset size.

## Footnotes

1 See the “Data Splits” and “Sequence Similarity” sections in (WPT) for example Preprint. Under review.

2 For example, it has been shown that Falcon-7B can be finetuned with a single 24GB NVidia RTX 4090

## Notes

https://huggingface.co/collections/AmelieSchreiber/esmbind-esmb-for-protein-binding-sites-64f9e4c3f210b74ac92b8b55

https://huggingface.co/collections/AmelieSchreiber/qbind-qlora-for-esm-2-binding-sites-prediction-651612ea01d873dd19124bfa

https://huggingface.co/collections/AmelieSchreiber/esm-ptm-esm-2-for-predicting-ptm-65206f49706c75514869aa96

### Competing Interest Statement

The authors have declared no competing interest.

